# Unveiling the crucial role of type IV secretion system and motility of *Helicobacter pylori* in IL-1β production via NLRP3 inflammasome activation in neutrophils

**DOI:** 10.1101/733790

**Authors:** Ah-Ra Jang, Min-Jung Kang, Jeong-Ih Shin, Soon-Wook Kwon, Ji-Yeon Park, Jae-Hun Ahn, Tae-Sung Lee, Dong-Yeon Kim, Bo-Gwon Choi, Myoung-Won Seo, Soo-Jin Yang, Min-Kyoung Shin, Jong-Hwan Park

**Author notes:** Address correspondence to: Min-Kyoung Shin,; Jong-Hwan Park.

## Abstract

*Helicobacter pylori* is a gram-negative, microaerophilic, and spiral-shaped bacterium and causes gastrointestinal diseases in human. IL-1β is a representative cytokine produced in innate immune cells and is considered to be a key factor in the development of gastrointestinal malignancies. However, the mechanism of IL-1β production by neutrophils during *H. pylori* infection is still unknown. We designed this study to identify host and bacterial factors involved in regulation of *H. pylori*-induced IL-1β production in neutrophils. We found that *H. pylori*-induced IL-1β production is abolished in NLRP3-, ASC-, and caspase-1/11-deficient neutrophils, suggesting essential role for NLRP3 inflammasome in IL-1β response against *H. pylori*. Host TLR2, but not TLR4 and Nod2, was also required for transcription of NLRP3 and IL-1β as well as secretion of IL-1β. *H. pylori* lacking *cagL*, a key component of the type IV secretion system (T4SS), induced less IL-1β production in neutrophils than did its isogenic WT strain, whereas *vacA* and *ureA* were dispensable. Moreover, T4SS was involved in caspase-1 activation and IL-1β maturation in *H. pylori*-infected neutrophils. We also found that FlaA is essential for *H. pylori*-mediated IL-1β production in neutrophils, but not dendritic cells. TLR5 and NLRC4 were not required for *H. pylori*-induced IL-1β production in neutrophils. Instead, bacterial motility is essential for the production of IL-1β in response to *H. pylori*. In conclusion, our study shows that host TLR2 and NLRP3 inflammasome and bacterial T4SS and motility are essential factors for IL-1β production by neutrophils in response to *H. pylori*.

**IMPORTANCE:** IL-1β is a representative pro-inflammatory cytokine and is considered to be a central host factor for the development of gastric cancers. Although neutrophils have been considered to be involved in *H. pylori*-induced gastric inflammation, the underlying mechanism by which *H. pylori* triggers IL-1β production in neutrophils remains to be defined. In this study, our data suggested a critical role for the host TLR2 and NLRP3 inflammasome in IL-1β production by neutrophil during *H. pylori* infection. Moreover, we found the bacterial factors, T4SS and FlaA, to be essential for IL-1β production and NLRP3 activation during the course of *H. pylori* infection. Our current findings provide detailed molecular genetic mechanisms associated with IL-1β production in neutrophils in response to *H. pylori* infection, which can serve as innovative anti-inflammatory targets to reduce *H. pylori*-induced gastric malignancies.

## INTRODUCTION

*Helicobacter pylori* (*H. pylori*) is a gram-negative, microaerophilic, and spiral bacterium that colonizes the human gastric mucosa. More than 50% of the world’s population are infected with the bacteria and the infection lasts a lifetime. *H. pylori* is the etiologic agent of gastrointestinal disorders that cause chronic gastritis, peptic ulcer, gastric adenocarcinoma, and gastric mucosa-associated lymphoid tissue (MALT) lymphoma (1). For this reason, it was classified by the World Health Organization as a class I carcinogen in 1994 (2).

Interleukin 1β (IL-1β) is considered to be a key factor correlated with development of gastric malignancies (3). Recently, polymorphisms of the *IL-1B* gene and IL-1 receptor antagonist (IL-1RN) have been revealed to be associated with *H. pylori*-related gastric cancer in the Chinese, Italian, and Indian population (4–6). Moreover, when infected with *H. felis*, IL-β-overexpressed transgenic mice display accelerated gastric inflammation development, metaplasia, and carcinoma (7). Huang *et al*. also showed that *H. pylori*-induced gastric inflammation and DNA methylation were reduced in IL-1R-deficient mice or by administration of IL-1 receptor antagonist (IL-1ra) (8). In addition, infiltration of innate immune cells, such as neutrophils and macrophages, and a multiplicity of gastric cancer induced by *H. pylori* infection were significantly reduced in IL-1β-deficient mice (9). Increased expression of IL-8, IL-1β, and COX-2 genes was also observed in patients with chronic gastritis infected with *H. pylori* compared with *H. pylori* negative patients (10). These findings suggest that IL-1β may play a crucial role in the development of *H. pylori*-induced gastric inflammation and cancer.

As a cytosolic multiprotein complex, NLRP3 inflammasome is a major inflammatory pathway that is activated in response to a variety of signals, including microbial infection and tissue damage (11). Activation of NLRP3 inflammasome is composed of two-step signals. First, the priming step is initiated by transcription of pro-IL-1β and NLRP3 through NF-κB activation by pattern-recognition receptors (PRRs) in response to microbial stimuli. The second step involves NLRP3 inflammasome oligomerization, caspase-1 activation, cleavage of pro-IL-1β by caspase-1, and then secretion of the mature IL-1β. This step is induced by various molecules, such as ATP, reactive oxygen species (ROS), and monosodium urate (MSU) (12). Several studies have demonstrated that *H. pylori* activates the NLRP3 inflammasome in innate immune cells, including dendritic cells (DCs) and neutrophils (13–16).

Neutrophils comprise about 50-70% of all leucocytes. These cells build a first line of defense against bacterial and fungal pathogens (17, 18). Despite the crucial role in innate immune response, several studies have reported that neutrophils might be involved in gastric-cancer development (19, 20). This concept has been supported by the observation that there was more neutrophils recruitment in gastric-cancer tissue than in the tissues surrounding gastric cancer (21). Furthermore, the higher number of neutrophils in gastric cancer is correlated with increased levels of IL-8 (21). In addition to the potential role of neutrophils in gastric-cancer development, a recent study has also shown that *H. pylori* T4SS induced production of IL-1β in human neutrophils in a NLRP3 inflammasome-dependent manner (15). However, the exact molecular mechanisms by which *H. pylori* bacterial factors regulate production of IL-1β in host neutrophils are not well defined. Thus, in this study, we sought to identify both bacterial and host factors associated with IL-1β production in neutrophils in response to *H. pylori* infection.

## RESULTS

### *H. pylori* induces IL-1β production in neutrophils by activating NLRP3 inflammasome by stimulating various danger signals

NLRP3 inflammasome has been known to be required for caspase-1 activation and IL-1β maturation in DCs in response to *H. pylori* (13, 14, 16). Therefore, we first investigated whether NLRP3 inflammasome is also involved in *H. pylori*-induced IL-1β maturation in neutrophils. As shown in Fig 1A, *H. pylori* P1WT induced IL-1β secretion in peritoneal neutrophils isolated from WT mice, whereas the production of IL-1β was abolished in NLRP3-, caspase-1/11-, and ASC-deficient neutrophils. Unlike the induction of IL-1β, *H. pylori*-induced TNF-α production was not affected by the same genetic deficiencies (Fig 1B). To confirm these data in a human-cell system, HL-60 cells, a human promyelocytic leukemia cell line, was differentiated to neutrophil-like cells at the presence of DMSO as previously described (22). As expected, *H. pylori* P1WT could induce IL-1β production in the differentiated HL-60 cells, which was reduced by the presence of glyburide (a NLRP3 inflammasome inhibitor) or Ac-YVAD-CMK (a caspase-1 inhibitor) in a dose-dependent manner (Fig 1C and D). Next, we did Western-blot analysis to detect cleaved IL-1β as a mature form. *H. pylori* P1WT led to cleavage of IL-1β in WT neutrophils, but not in cells from NLRP3-, caspase-1/11-, and ASC-deficient mice (Fig 1E). These findings suggest that the NLRP3/ASC/caspase-1 axis is essential for *H. pylori*-induced IL-1β production in neutrophils.

**Figure 1.**
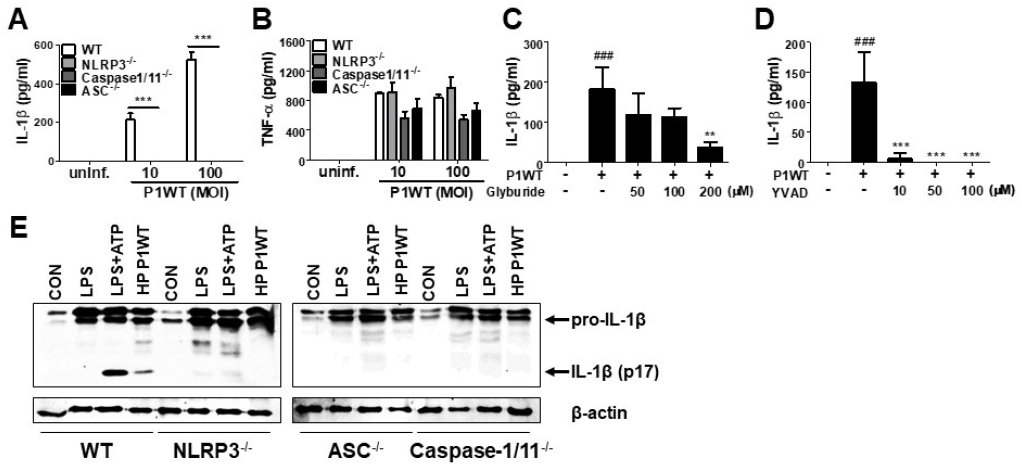
*H. pylori* mediates production of IL-1β in NLRP3 in an inflammasome-dependent manner in neutrophils. WT, and NLRP3-, Caspase1/11-, and ASC-deficient mouse peritoneal neutrophils were infected with *H. pylori* P1WT (MOI 100) for 24 h (A and B). The concentration of TNF-α and IL-1β in supernatant was measured by ELISA. HL-60 cells were pretreated without or with glyburide (C) or Ac-YVAD-CMK (D) at the indicated concentrations for 2 h and subsequently infected with *H. pylori* P1WT (MOI 100) for 24 h. The concentration of IL-1β in supernatant was measured by ELISA. We infected peritoneal neutrophils from WT and NLRP3-, ASC-, and Caspase1/11-deficient mice with *H. pylori* P1WT (MOI 100) for 6 h (E). We used culture supernatants and cell lysates to detect immature and cleaved forms of caspase-1 and IL-1β by Immunoblotting. Antibody against β-actin was used as a loading control. Results are presented as mean ± SD. ###, *p* < 0.001 *vs*. control cells. **, *p* < 0.01; ***, *p* < 0.001 *vs.* HP P1WT infected with cells.

Various signals, such as extracellular ATP, K^+^ efflux, reactive oxygen species (ROS) generation, and cathepsin B release from lysosomes, mediates activation of NLRP3 inflammasome (12). To find out whether these stimuli are involved in *H. pylori*-induced IL-1β production in neutrophils, we carried out inhibitor assays. Oxidized ATP (oxATP) (a P2X7R antagonist) and extracellular addition of KCl reduced IL-1β production in *H. pylori* P1WT-infected neutrophils in a dose-dependent manner, and the cytokine production was completely abolished at a high concentration of those inhibitors, whereas TNF-α level was slightly influenced (Fig S1A-D). A ROS inhibitor NAC reduced both IL-1β and TNF-α at 20 mM concentration (Fig S1E and F). The IL-1β production was significantly inhibited by CA074Me, a cathepsin B inhibitor, even at a low concentration of 5 μM, which did not influence TNF-α level (Fig S1G and H). These findings suggest that various extracellular signals, such as ATP/P2X7R, potassium efflux, ROS generation, and cathepsin B, may contribute to *H. pylori*-induced activation of the NLRP3 inflammasome pathway in neutrophils.

### TLR2, but not Nod2 and TLR4, is required for expression of IL-1β gene and secretion of IL-1β in *H. pylori*-infected neutrophils

Previously, TLR2 and Nod2 have been shown to be involved in *H. pylori*-mediated gene expression of NLRP3 and IL-1β in DCs (13). Moreover, *H. pylori* upregulates TLR4 expression in human gastric epithelial cells (23). We therefore explored whether such patterns of recognition mediate IL-1β production in neutrophils in response to *H. pylori* infection. TLR2 deficiency led to impaired production of both TNF-α and IL-1β in *H. pylori* P1WT-infected peritoneal neutrophils derived from mice (Fig. 2A and B). In contrast, Nod2 and TLR4-deficient cells produced a comparable level of IL-1β and TNF-α (Fig 2A and B). Consistently, *H. pylori*-induced production of IL-1β and TNF-α was impaired in TLR2-deficient bone marrow-derived neutrophils (BMDNs) (Fig. 2C and D). To find out whether TLR2 contributes to priming of NLRP3 and IL-1β (signal 1), we evaluated their gene expression by RT-PCR. As shown by the results in Fig. 2E-H, we observed increased expression of genes for NLRP3 and IL-1β in WT peritoneal neutrophils and BMDNs, whereas we observed significantly lower levels of NLRP3 and IL-1β gene transcription in TLR2-deficient cells (Fig. 2E-H). Collectively, these data indicate that TLR2 is a central innate receptor for regulating NLRP3 inflammasome priming (signal) in *H. pylori*-infected neutrophils.

**Figure 2.**
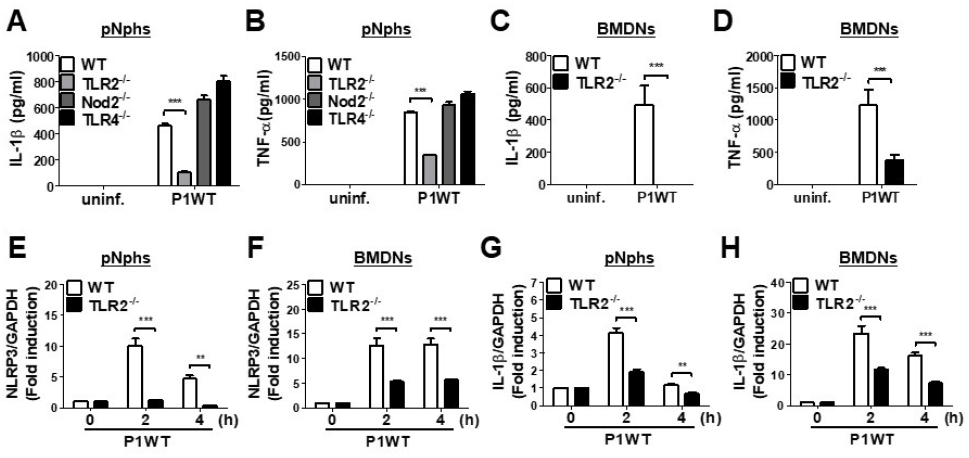
TLR2, but not NOD2 and TLR4, is involved in *H. pylori* recognition in mouse neutrophils. WT and TLR2-, NOD2-, and TLR4-deficient peritoneal neutrophils (A and B) and BMDNs (C and D) from WT and TLR2-deficient mice were infected with P1WT (MOI 100) for 24 h. The concentration of IL-1β (A and C) and TNF-α (B and D) in the supernatant was measured by ELISA. We infected WT and TLR2-deficient peritoneal neutrophils or BMDNs with *H. pylori* P1WT (MOI 100) at indicated times (E-H). Gene expressions of NLRP3 (E and F) and IL-1β (G and H) were evaluated by real-time PCR. Results are presented as mean ± SD. **, *p* < 0.01; ***, *p* < 0.001.

### *H. pylori* Type IV secretion system is required for IL-1β secretion through caspase-1 activation and IL-1β processing in neutrophils

A previous report suggested that *H. pylori* uses a type IV secretion system (T4SS) to induce pro-IL-1β expression in DCs (13). In contrast, a recent study indicated that the *H. pylori* T4SS is dispensable for IL-1β production in human neutrophils (15). To assess the role of T4SS in production of IL-1β in various immune cells, we infected peritoneal neutrophils, BMDNs, and the differentiated HL-60 cells with *H. pylori* P1WT and its isogenic mutant P1*ΔCagL*, in which T4SS is non-functional (24). As expected, we observed impaired IL-1β production in peritoneal neutrophils infected with P1*ΔcagL* (Fig 3A). Consistently, *cagL* deficiency reduced the production of IL-1β in both BMDNs and human neutrophil-like HL-60 cells (Fig 3B and C). In contrast to the production of IL-1β, the *cagL* mutant induced a comparable level of TNF-α in each cell type to its isogenic WT strain (Fig 3D-F). Western blot analysis revealed that cleavage of caspase-1 and IL-1β was reduced in P1*ΔCagL*-treated neutrophils more than in cells infected with P1WT (Fig 3G). There was no difference in expression level of the pro-form of IL-1β and casapse-1 between P1WT- and *ΔCagL*-infected neutrophils (Fig S2). In addition, cagL deficiency did not influence the gene expression of NLRP3 and IL-1β in either peritoneal neutrophils or BMDNs (Fig S3). These findings demonstrated that bacterial T4SS is involved in regulation of NLRP3 inflammasome activation (signal 2) rather than priming (signal 1) in host neutrophils in response to *H. pylori* infection.

**Figure 3.**
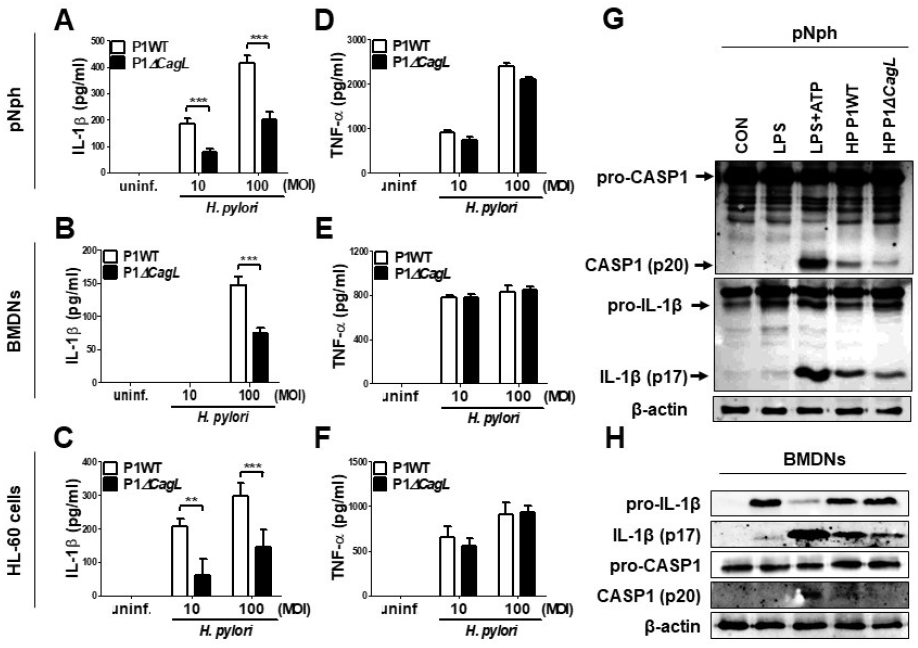
*H. pylori* T4SS induces production of IL-1β in neutrophils. Peritoneal neutrophils (A and D), BMDNs (B and E), and HL-60 cells (C and F) were infected with *H. pylori* P1WT and isogenic mutant deficient in cagL (MOI 10 and 100) for 24 h. The concentration of IL-1β (A-C) and TNF-α (D-F) in supernatant was measured by ELISA. Results are presented as mean ± SD. **, *p* < 0.01; ***, *p* < 0.001. Peritoneal neutrophils (G) and BMDNs (H) were infected with P1WT and ∆*cagL* (MOI 100) for 6 h. We used culture supernatants and cell lysates to detect immature and cleaved forms of caspase-1 and IL-1β by Immunoblotting (G and H). Antibody against β-actin was used as a loading control.

### Deficiency of FlaA, but not of UreA and VacA, distinctly leads to impaired IL-1β secretion in *H. pylori*-infected neutrophils

It has been reported that the *H. pylori* virulence factors, such as VacA and UreA, are involved in IL-1β production and NLRP3 inflammasome activation in DCs (14, 16). Bacterial flagellin has also been proposed to be involved in IL-1β processing via NLRC4 inflammasome activation (25). Therefore, we investigated whether such bacterial factors are required for *H. pylori*-induced IL-1β production in neutrophils. As shown in Fig. 4A and B, both UreA and VacA mutant strains exhibited a similar level of IL-1β to the isogenic WT strain in peritoneal neutrophils, whereas TNF-α induction was slightly decreased in the two mutant strains (Fig 4A and B). Unexpectedly, IL-1β production was significantly lower in P1*∆FlaA*-treated neutrophils than in cells infected with the P1WT strain, whereas levels of TNF-α were comparable between the two strains (Fig 4A and B). In BMDNs, the FlaA deficiency also impaired the production of IL-1β, but not TNF-α (Fig. 4C and D). Neither UreA nor VacA affected IL-1β and TNF-α production in BMDNs in response to *H. pylori* (Fig, 4C and D). To confirm these data in a different *H. pylori* genetic background strain, we infected peritoneal neutrophils with *H. pylori* 26695WT and its isogenic mutant 26695*∆FlaA*. In agreement with the data generated with the P1WT strain, IL-1β level was significantly lower in 26695*∆FlaA*-treated cells than in ones treated with the 26695WT strain, whereas the TNF-α level was not affected by deletion of the FlaA gene (Fig 4E and F). To find out whether FlaA is required for IL-1β production in DCs, we infected BMDCs with *H. pylori* P1WT and P1*∆FlaA*. As shown in Fig. 4G, the P1WT and P1*∆FlaA* strains displayed comparable levels of IL-1β secretion in DCs. These results indicated that *H. pylori* FlaA distinctly contributes to production of IL-1β in neutrophils, but not in DC.

**Figure 4.**
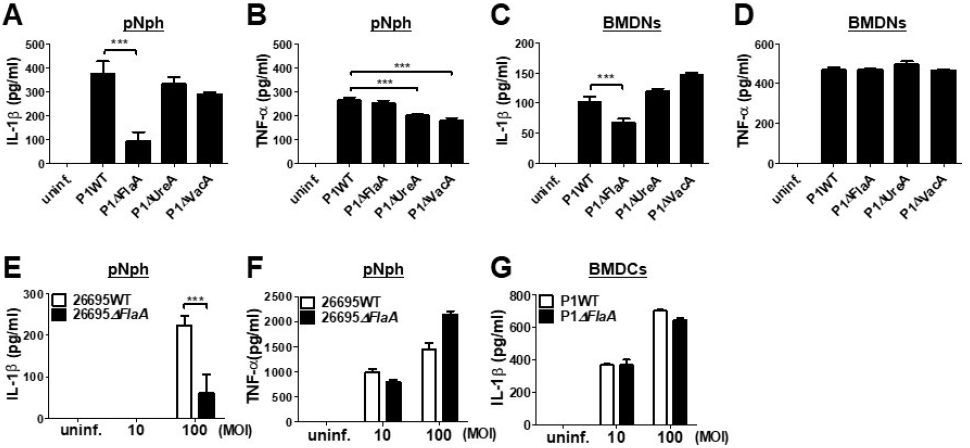
*H. pylori* flagellin is important for IL-1β production in mouse neutrophils, but not BMDCs. Peritoneal neutrophils (A, B, E, and F), BMDNs (C and D), and BMDCs (G) were infected with indicated bacteria (MOI 100) for 24 h. We measured the concentration of IL-1β (A, C, E, and G) and TNF-α (B, D, and F) in the supernatant by ELISA. Results are presented as mean ± SD. ***, *p* < 0.001

### FlaA contributes to both priming and inflammasome activation of NLRP3 in neutrophils in response to *H. pylori*

We next explored whether FlaA-induced production of IL-1β is mediated via activation of NLRP3 inflammasome in neutrophils. *H. pylori* P1WT induced strong expression of pro-IL-1β protein, which was slightly reduced in the cells treated with P1*∆FlaA* (Fig 5A). P1WT-infected BMDNs also released a significantly higher level of cleaved IL-1β (mature form) than did BMDNs infected with the P1*∆FlaA* strain (Fig. 5A). Similar to the P1*∆FlaA* strain, FlaA deficiency in the 26695 background also led to reduced cleavage of IL-1β and thus to less pro IL-1β expression (Fig 5B). In addition, expression of NLRP3 and IL-1β genes was significantly lower in P1*∆FlaA*-treated BMDNs P1WT-treated cells (Fig. 5E and F). Consistently, NLRP3 gene expression was reduced in BMDNs in response to FlaA deficiency in the *H. pylori* 26695 strain (Fig 5G). Moreover, IL-1β expression was also decreased in 26695*∆FlaA*-treated cells at 2 h after infection (Fig 5H). These data indicate that *H. pylori* flagellin is involved in regulation of both priming and activation of NLRP3 inflammasome in neutrophils.

**Figure 5.**
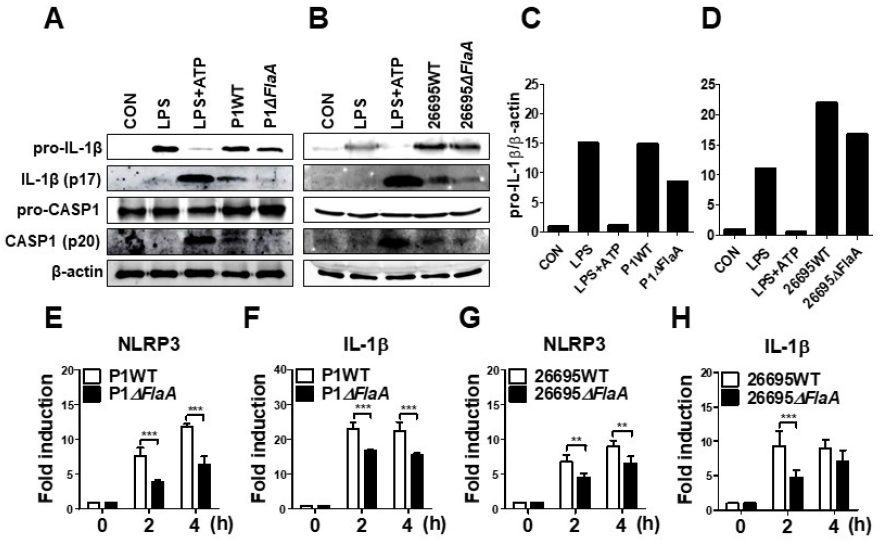
*H. pylori* flagellin induces activation of caspase-1 and IL-1β in response to *H. pylori* in mouse neutrophils. BMDNs were infectedwith *H. pylori* P1WT and ∆*flaA* (A and C) or 26695WT and ∆*flaA* (B and D) (MOI 100) for 6 h. We used culture supernatants and cell lysates to detect immature and cleaved forms of caspase-1 and IL-1β by Immunoblotting (A-D). Antibody against β-actin was used as a loading control. Peritoneal neutrophils (E and F) and BMDNs (G and H) were infected with *H. pylori* P1WT (MOI 100) at indicated time. We evaluated gene expression of NLRP3 (E and G) and IL-1β (F and H) by real-time PCR. Results are presented as mean ± SD. **, *p* < 0.01; ***, *p* < 0.001.

### Bacterial motility, but not host TLR5 and NLRC4, is required for *H. pylori*-mediated IL-1β production in neutrophils

TLR5 and NLRC4 are dual sensors for bacterial flagellin at the cell surface and cytosol in host cells (24). Flagellin recognition by TLR5 leads to NF-κB activation, whereas NLRC4 is responsible for inflammasome activation. Since *H. pylori* FlaA was essential for IL-1β gene expression and maturation in neutrophils, we tried to find out whether TLR5 and NLRC4 are involved in *H. pylori*-mediated IL-1β production. As shown by the results in Fig. 6A and B, WT neutrophils and neutrophils deficient of TLR5 or NLRC4 produced a comparable level of IL-1β and TNF-α in response to *H. pylori* (Fig. 6A and B), suggesting that *H. pylori* FlaA regulates IL-1β production in neutrophils in a TLR5-and NLRC4-independent manner. Because bacterial flagella are essential for motility, we next explored whether motility is essential for *H. pylori*-induced IL-1β production in neutrophils. In our *in vitro* system, we cultured neutrophils in a floating state. Therefore, we centrifuged the cells after bacterial infection to increase the contact between *H. pylori* and neutrophils. As shown in Fig, 6C, the centrifuging abolished the difference of IL-1β production by WT and *ΔFlaA* mutant, suggesting that motility may be involved in *H. pylori*-induced IL-1β production in neutrophils. We further investigated whether different levels of motility in clinical isolates of *H. pylori* correlate with the amount of IL-1β production in neutrophils. Five clinical isolates of *H. pylori* were divided into two groups based on the results from a bacterial motility assay (Fig. S4): a low motility group of two strains (4940A and 4980AC) and a high motility group with three strains (4930AC, 5049AC, and 5356AC). *H. pylori* strains with high motility induced increased levels of IL-1β in neutrophils versus the strains with low motility (Fig. 6D), whereas TNF-α level was not different between the two groups of strains (Fig. 6E). Furthermore, *H. pylori* 52 strain originally displayed a low motility phenotype, but it obtained high motility phenotype (HP 52 P6) after six passages in mice (data not shown). Interestingly, the passaged *H. pylori* strain induced a significantly higher level of IL-1β in neutrophils than did the original strain with low motility (Fig 6F). The two strains did differ in induction of TNF-α production (Fig 6G).

**Figure 6.**
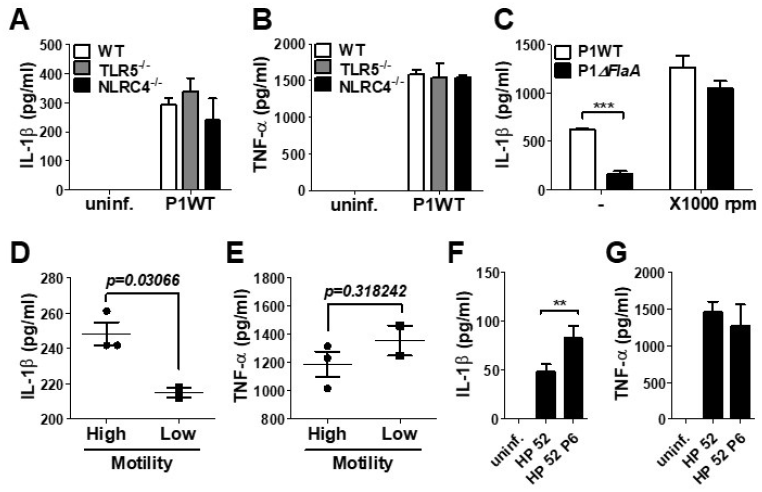
The production of IL-1β is regulated by *H. pylori* motility of flagellin, but not NLRC4 and TLR5 in mouse neutrophils. WT and NLRC4-, and TLR5-deficient peritoneal neutrophils were infected with *H. pylori* P1WT (MOI 100) for 24 h (A and B). To promote the contact between cells and *H. pylori*, we used mild centrifuging (1000 rpm, 10 min). Clinical isolates of *H. pylori* with or without motility were infected in peritoneal neutrophils (D and E). *H. pylori* 52 and mouse-adapted bacteria (passage 6) were infected in peritoneal neutrophils (F and G). The concentration of IL-1β (A, C, D, and F) and TNF-α (B, E, and G) in supernatant was measured by ELISA. Results are presented as mean ± SD. **, *p* < 0.01, two-tailed Student’s *t*-test.

## DISCUSSION

IL-1β is considered to be a central factor for gastric malignancy, as is supported by evidence that *IL-1B* gene polymorphism increases the risk of gastric cancer (26, 27) and overexpression of IL-1β leads to gastric inflammation and cancers in mice (7, 9). It has been known that *H. pylori* leads to IL-1β production in DCs through an NLRP3-dependent pathway (13, 14, 16). Neutrophils are key innate immune cells that are involved in *H. pylori*-mediated gastric inflammation (28). Therefore, it is conceivable that both bacterial and host factors may play crucial roles in the production of IL-1β in neutrophils. However, only limited information is available on the precise molecular mechanisms involved in *H. pylori*-induced IL-1β production in neutrophils.

Recently, Pérez-Figueroa *et al*. reported that *H. pylori* induces IL-1β production in human neutrophils in a TLR2- and TLR4-independent manner (15). They also demonstrated that *H. pylori* increases the expression of NLRP3 inflammasome components and inhibitors for NLRP3, and caspase-1 reduces the production of IL-1β (15). Moreover, in contrast to the important role of T4SS in DCs, T4SS was dispensable for *H. pylori*-induced IL-1β production in human neutrophils (15). However, some aspects are inconsistent with our current study. In this study, we identified that NLRP3 inflammasome is a key host factor in neutrophils for production of IL-1β in response to *H. pylori* by showing that secretion and cleavage of IL-1β were abolished in NLRP3-, caspase-1/11-, and ASC-deficient neutrophils. However, unlike the previous report by Pérez-Figueroa *et al*. (15), our data suggested that TLR2 is required for *H. pylori*-induced IL-1β production in both peritoneal neutrophils and BMDNs. Expression of NLRP3 and IL-1β genes was also reduced in TLR2-deficient neutrophils. In *H. pylori*-infected DCs, TLR2 has been known to mediate IL-1β production by regulating NLRP3 and pro IL-1β expression (signal 1) (13, 14). Taken together, it is likely that TLR2 contributes to IL-1β production in response to *H. pylori* by activating expression of NLRP3 and IL-1β genes in innate immune cells, which include neutrophils. In addition, it seems to be necessary to measure the induction levels of NLRP3 or IL-1β expression in the human neutrophils prepared in the reports by Pérez-Figueroa *et al*., because TLR2 may be dispensable if the cells are primed sufficiently.

Nod2 is a representative cytosolic pattern-recognition receptor that is responsible for the recognition of a bacterial peptidoglycan motif and muramyl dipeptide (MDP). After MDP recognition, Nod2 interacts with its adaptor protein receptor interacting protein kinase 2 (Ripk2) through CARD-CARD interaction, followed by activation of NF-κB and mitogen activated protein kinases (MAPKs) (24). Nod2 is known to be functionally expressed in human and murine neutrophils (29, 30). However, we do not know the precise mechanism by which Nod2 is involved in the recognition of *H. pylori* in immune cells including neutrophils. Kim *et al*. showed that gene and protein expression of IL-1β in response to *H. pylori* are reduced in Nod2-deficient BMDCs as compared to WT cells (13). In contrast, in a study by Koch *et al*., the authors claimed that Nod2 does not contribute to *H. pylori*-mediated caspase-1 activation and IL-1β production in BMDCs. Currently, it is unclear whether this discrepancy results from differences in bacterial strains used and/or in experimental condition. In macrophages, microarray analysis revealed that there was no substantial changes of gene expression between MyD88/TRIF- and MyD88/TRIF/Ripk2-deficient cells during *H. pylori* infection (31), suggesting that the Nod1/Nod2 signaling pathway may not be important in immune response of macrophages against *H. pylori*. In the present study, the IL-1β level induced by *H. pylori* was comparable between WT and Nod2-deficient neutrophils. Therefore, it is likely that Nod2 does not play a critical role in neutrophils’ function against *H. pylori*.

Several bacterial factors including T4SS, vacuolating toxin, and urease contribute to IL-1β production in response to *H. pylori* in BMDCs (13, 14, 16). Isogenic mutants lacking both *ureA* and *ureB* genes failed to activate caspase-1, whereas transcription of IL-1β was unaffected (14). The *vacA* mutant also led to less production of IL-1β and impaired caspase-1 activation in LPS-primed BMDCs (16). There is still a controversy on the precise role of T4SS in regulation of IL-1β production in DCs. Kim *et al*. reported that the cagL mutant produced level of IL-1β similar to that of the isogenic WT strain in LPS-primed DCs, in which NLRP3 and IL-1β are sufficiently induced, whereas mRNA expression of IL-1β and its secretion were reduced in cagL mutant-treated cells in the unprimed condition (13), suggesting that T4SS is essential for the priming step. In contrast, Semper *et al*. showed that a mutant lacking cagE produced less IL-1β in LPS-primed DCs (16). In the present study, we demonstrated that T4SS, but not VacA and UreA, is required for *H. pylori*-induced IL-1β production in neutrophils. In *H. pylori*-infected neutrophils, T4SS seems to regulate signal 2 of inflammasome activation rather than priming of NLRP3 and pro IL-1β, because cagL deficiency did not influence protein expression of pro IL-1β or gene expression of NLRP3 and IL-1β. Our current results are inconsistent with the previous report that suggested a dispensable role of T4SS in *H. pylori*-induced production of IL-1β (15). In the previous study, the experimental design apparently was flawed by using the *H. pylori* 26695 strain as the WT strain and the virD4 mutant originating from a different genetic background, *H. pylori* G27 strain (15). Because the ability to produce IL-1β can differ among bacterial strains, this should be confirmed by using isogenic set of *H. pylori* strains.

It is remarkable that the FlaA mutant led to less production of IL-1β in response to *H. pylori* in neutrophils, as was confirmed by using two sets of *H. pylori* strains, P1 and 26695. This seems to be specific in neutrophils, because FlaA deficiency did not influence IL-1β production in BMDCs. In addition, we showed that FlaA was required for gene transcription of NLRP3 and IL-1β and cleavage of caspase-1. TLR5 and NLRC4 are two central host receptors for the recognition of bacterial flagellin. TLR5 sensing of flagellin triggers NF-κB activation, followed by the production of pro-inflammatory cytokines (32). Bacterial flagellin can induce IL-1β production in innate immune cells through a NAIP-NLRC4-dependent pathway (33). However, in this study, both TLR5- and NLRC4-deficient neutrophils could produce a comparable level of IL-1β in response to *H. pylori*. In fact, it is known that *H. pylori* FlaA leads to weak activation of TLR5, and its purified flagellin fails to induce IL-8 production and p38 MAPK activation in gastric epithelial cells (34). Moreover, in contrast to *Salmonella Typhimurium*, *H. pylori* flagellin could not induce activation of caspase-1 and production of IL-18 and LDH in macrophages, although it induced NLRC4 Ser533 phosphorylation (35). These findings indicate that *H. pylori* flagellin may contribute to production of IL-1β in neutrophils via a TLR5- and NLRC4-independent pathway. Instead, our results suggested that bacterial motility is essential for the production of IL-1β in response to *H. pylori*, as is supported by evidence that centrifuging that enhances bacteria to cell contact abolished the difference of IL-1β level produced by *H. pylori* P1WT and isogenic FlaA mutants and that clinical isolates with high motility produced more IL-1β than those with low motility. SINCE this phenomenon was seen only in neutrophils, but not BMDCs, a further study should be done to clarify the cell-type specific role of *H. pylori* flagellin in production of IL-1β.

In conclusion, we shows here that *H. pylori* T4SS and flagellin are essential for IL-1β production in neutrophils. TLR2 and NLRP3 inflammasome are central host factors to regulate neutrophil production of IL-1β in response to *H. pylori*. Since IL-1β plays an important role in the development of gastric malignancies, these bacterial and host factors can be preventive and therapeutic targets for IL-1β-mediated gastric abnormalities.

## MATERIALS AND METHODS

### Mice

We purchased wild type (WT), TLR2-, TLR4-, and NOD2-deficient mice on C57BL/6 background from the Jackson Laboratory (Bar Harbor, ME, USA). NLRP3-, Capase-1/11-, ASC-, and NLRC4-deficient mice were kindly provided by Prof. Gabriel Núñez (University of Michigan, USA). TLR5-deficient mice were gifts from Prof. Joon Haeng Rhee (Chonnam National University, Hwasun, Korea). We conducted all animal studies using protocols approved by the Institutional Animal Care and Use Committee of Chonnam National University (Approval No. CNU IACUC-YB-2018-85).

### Bacterial strains and culture conditions

*H. pylori* P1WT and its isogenic mutants P1∆*CagL*, P1∆*FlaA*, P1∆*UreA*, and P1*∆VacA* have been described previously (36). Another mutant with FlaA deficiency was generated by allelic exchange in *H. pylori* 26695 strain, and details are provided in the Supplemental Material. The following clinical isolates from child patients were provided from Pyeongyang National University Hospital (GNUH), as the Branch of National Culture Collection for Pathogens (NCCP, Jinju, Korea): three motile strains, *H. pylori* 5356AC*, H. pylori* 4930AC*, H. pylori* 5049AC; two non-motile strains, *H. pylori* 4940A*, H. pylori* 4980AC. *H. pylori* 52WT (non-motile) and its mouse-adapted strain *H. pylori* 52P6 (six time-passaged) were also provided from Pyeongyang National University Hospital. We cultured all *H. pylori* strains on Brucella broth containing 10% fetal bovine serum (FBS; Corning costar, Corning NY, USA), 1 μg/ml nystatin (Sigma-Aldrich, St. Louis, MO, USA), 5 μg/ml trimethoprim (Sigma-Aldrich), and 10 μg/ml vancomycin (Sigma-Aldrich) at 37°C under microaerobic conditions.

### Reagent and inhibitor assay

We purchased LPS from *Escherichia coli* O111:B4 from InvivoGen (San Diego, CA, USA). Adenosine 5′-triphosphate disodium salt hydrate (ATP) was purchased from Sigma-Aldrich. For inhibitor assay, we used glyburide (Sigma-Aldrich) as inhibition of NLRP3 and Ac-YVAD-CMK (Calbiochem, La. Jolla, CA, USA) as inhibition of Caspase-1.

### Cell culture

We isolated thioglycollate-induced peritoneal neutrophils as previously described (37). Briefly, mouse peritoneal neutrophils were harvested 4 h after intraperitoneal injection of 2 ml of 4% thioglycollate broth (Sigma-Aldrich). The collected peritoneal neutrophils were cultured in RPMI 1640 (Welgene, Gyeongsan, Gyeongsangbuk-do, Korea) containing 10% FBS in a 5% CO_2_ incubator at 37°C. To obtain of neutrophils derived from bone marrow, we isolated cells from femurs and tibias using density gradient cell separation protocol. Total bone marrow cells were overlaid on a two-layer gradient of HISTOPAQUE-1119 (density: 1.119 g/ml; Sigma-Aldrich) and HISTOPAQUE-1077 (density: 1.077 g/ml; Sigma-Aldrich) and centrifuged (2000 rpm, 30 min) without braking. The collected cells in the interface were used. Bone-marrow-derived neutrophils (BMDNs) were resuspended in RPMI 1640 (Welgene) containing 10% FBS in a 5% CO_2_ incubator at 37°C. Human leukemia cell line HL-60 was cultured in RPMI 1640 medium (Welgene) containing 10% FBS, 1% Penicillin/Streptomycin (P/S; Gibco, Grand Island, NY, USA), 2 mM L-glutamine (Gibco), and 25 mM HEPES (Gibco). To differentiate into neutrophil-like cells, we stimulated cells with 1.25% DMSO for seven days in a 5% CO_2_ incubator at 37°C. Bone marrow-derived dendritic cells (BMDCs) were isolated and differentiated as described previously (38). Briefly, BMDCs were cultured in RPMI 1640 (Welgene) containing 10% FBS, 1% P/S, 2 mM L-glutamine, 50 μM 2-mercaptoethanol (Sigma-Aldrich), and 20 ng/ml GM-CSF (Peprotech, Rocky Hill, NJ, USA) in a 5% CO2 incubator at 37°C for 9 days. Fresh medium was added both three and six days later. We seeded the cells in 6-well or 48-well plates at a density of 2×10^6^ cells or 2×10^5^ cells and incubated them in a 5% CO2 incubator at 37°C, and then infected them with *H. pylori*.

### Measurement of cytokines

We measured the concentrations of IL-1β and tumor necrosis factor alpha (TNF-α) in culture supernatants by using a commercial ELISA kit (R&D Systems, Minneapolis, MN, USA).

### Immunoblotting

Culture supernatants and remaining cells were lysed by 1% Triton-X 100 (Sigma-Aldrich) and a complete protease inhibitor cocktail (Roche, Mannheim, Germany). After centrifugation at 3000 rpm for 5 min, we mixed the supernatant with SDS-PAGE sample loading buffer (5×). To detect target proteins, samples were separated by 15% SDS-PAGE and transferred to nitrocellulose (NC) membranes. We probed membranes with primary antibodies against caspase-1 (Enzo Life Science, Farmingdale, NY, USA), IL-1β (R&D Systems), and β-actin (Santa Cruz Biotechnology, Dallas, TX, USA). After immunoblotting with secondary antibodies (Thermo Fisher Scientific, MA, USA), we detected proteins using Clarity Western ECL Substrate (Bio-Rad, Hercules, CA, USA).

### Real-time quantitative PCR (qPCR)

RNA was extracted using easy-BLUE^TM^ Total RNA Extraction Kit (Intron Biotechnology, Seongnam, Korea) and cDNA synthesis was done using ReverTra Ace^®^ qPCR RT Master Mix (TOYOBO Bio-Technology, Osaka, Japan) according to the manufacturer’s instructions. qPCR was performed by the CFX Connect^TM^ Real-time PCR Detection System (Bio-Rad, Hercules, CA, USA) using 2x PCRBIO SyGreen Blue Mix Lo-ROS according to the manufacturer’s instructions (Bio-D Co., Ltd,, Hull, UK). GAPDH was used for normalization. The primers used for qPCR were as follows:

mNLRP3 forward: 5’-ATGGTATGCCAGGAGGACAG-3’;

mNLRP3 reverse: 5’-ATGCTCCTTGACCAGTTGGA-3’;

mIL-1β forward: 5’-GATCCACACTCTCCAGCTGCA-3’;

mIL-1β reverse: 5’-CAACCAACAAGTGATATTCTCCATG-3’;

mGAPDH forward: 5’-CGACTTCAACAGCAACTCCCACTCTTCC-3’;

mGAPDH reverse: 5’-TGGGTGGTCCAGGGTTTCTTACTCCTT-3’.

### Statistical analysis

Statistical significance of differences among groups was found by using two-tailed Student’s *t*-test or the one-or two-way analysis of variance (ANOVA) followed by Bonferroni post-tests. We calculated all statistics using GraphPad Prism version 5.01 (GraphPad Software, San Diego, CA, USA). Values of *p* < 0.05 were considered statistically significant.

## ACKNOWLEDGMENTS

This research was supported by the Mid-Career Researcher Program (Grant No. 2018R1A2B3004143) of the National Research Foundation of Korea (NRF) funded by the Ministry of Science and ICT (Information and Communication Technologies). The funders had no role in the study design, data collection, or interpretation, or in the decision to submit the work for publication.

## REFERENCES

1. Kusters JG, van Vliet AH, Kuipers EJ. 2006. Pathogenesis of Helicobacter pylori infection. Clin Microbiol Rev 19:449–90.

2. Anonymous. 1994. Schistosomes, liver flukes and Helicobacter pylori. IARC Working Group on the Evaluation of Carcinogenic Risks to Humans. Lyon, 7-14 June 1994. IARC Monogr Eval Carcinog Risks Hum 61:1–241.

3. El-Omar EM, Carrington M, Chow WH, McColl KE, Bream JH, Young HA, Herrera J, Lissowska J, Yuan CC, Rothman N, Lanyon G, Martin M, Fraumeni JF, Jr., Rabkin CS. 2000. Interleukin-1 polymorphisms associated with increased risk of gastric cancer. Nature 404:398–402.

4. Yang J, Hu Z, Xu Y, Shen J, Niu J, Hu X, Guo J, Wei Q, Wang X, Shen H. 2004. Interleukin-1B gene promoter variants are associated with an increased risk of gastric cancer in a Chinese population. Cancer Lett 215:191–8.

5. Palli D, Saieva C, Luzzi I, Masala G, Topa S, Sera F, Gemma S, Zanna I, D’Errico M, Zini E, Guidotti S, Valeri A, Fabbrucci P, Moretti R, Testai E, del Giudice G, Ottini L, Matullo G, Dogliotti E, Gomez-Miguel MJ. 2005. Interleukin-1 gene polymorphisms and gastric cancer risk in a high-risk Italian population. Am J Gastroenterol 100:1941–8.

6. Kumar S, Kumar A, Dixit VK. 2009. Evidences showing association of interleukin-1B polymorphisms with increased risk of gastric cancer in an Indian population. Biochem Biophys Res Commun 387:456–60.

7. Tu S, Bhagat G, Cui G, Takaishi S, Kurt-Jones EA, Rickman B, Betz KS, Penz-Oesterreicher M, Bjorkdahl O, Fox JG, Wang TC. 2008. Overexpression of interleukin-1beta induces gastric inflammation and cancer and mobilizes myeloid-derived suppressor cells in mice. Cancer Cell 14:408–19.

8. Huang FY, Chan AO, Lo RC, Rashid A, Wong DK, Cho CH, Lai CL, Yuen MF. 2013. Characterization of interleukin-1beta in Helicobacter pylori-induced gastric inflammation and DNA methylation in interleukin-1 receptor type 1 knockout (IL-1R1(-/-)) mice. Eur J Cancer 49:2760–70.

9. Shigematsu Y, Niwa T, Rehnberg E, Toyoda T, Yoshida S, Mori A, Wakabayashi M, Iwakura Y, Ichinose M, Kim YJ, Ushijima T. 2013. Interleukin-1beta induced by Helicobacter pylori infection enhances mouse gastric carcinogenesis. Cancer Lett 340:141–7.

10. Bartchewsky W, Jr., Martini MR, Masiero M, Squassoni AC, Alvarez MC, Ladeira MS, Salvatore D, Trevisan M, Pedrazzoli J, Jr., Ribeiro ML. 2009. Effect of Helicobacter pylori infection on IL-8, IL-1beta and COX-2 expression in patients with chronic gastritis and gastric cancer. Scand J Gastroenterol 44:153–61.

11. Man SM, Kanneganti TD. 2015. Regulation of inflammasome activation. Immunol Rev 265:6–21.

12. Franchi L, Eigenbrod T, Munoz-Planillo R, Nunez G. 2009. The inflammasome: a caspase-1-activation platform that regulates immune responses and disease pathogenesis. Nat Immunol 10:241–7.

13. Kim DJ, Park JH, Franchi L, Backert S, Nunez G. 2013. The Cag pathogenicity island and interaction between TLR2/NOD2 and NLRP3 regulate IL-1beta production in Helicobacter pylori infected dendritic cells. Eur J Immunol 43:2650–8.

14. Koch KN, Hartung ML, Urban S, Kyburz A, Bahlmann AS, Lind J, Backert S, Taube C, Muller A. 2015. Helicobacter urease-induced activation of the TLR2/NLRP3/IL-18 axis protects against asthma. J Clin Invest 125:3297–302.

15. Perez-Figueroa E, Torres J, Sanchez-Zauco N, Contreras-Ramos A, Alvarez-Arellano L, Maldonado-Bernal C. 2016. Activation of NLRP3 inflammasome in human neutrophils by Helicobacter pylori infection. Innate Immun 22:103–12.

16. Semper RP, Mejias-Luque R, Gross C, Anderl F, Muller A, Vieth M, Busch DH, Prazeres da Costa C, Ruland J, Gross O, Gerhard M. 2014. Helicobacter pylori-induced IL-1beta secretion in innate immune cells is regulated by the NLRP3 inflammasome and requires the cag pathogenicity island. J Immunol 193:3566–76.

17. Freitas M, Lima JL, Fernandes E. 2009. Optical probes for detection and quantification of neutrophils’ oxidative burst. A review. Anal Chim Acta 649:8–23.

18. Malech HL, Deleo FR, Quinn MT. 2014. The role of neutrophils in the immune system: an overview. Methods Mol Biol 1124:3–10.

19. Zhao JJ, Pan K, Wang W, Chen JG, Wu YH, Lv L, Li JJ, Chen YB, Wang DD, Pan QZ, Li XD, Xia JC. 2012. The prognostic value of tumor-infiltrating neutrophils in gastric adenocarcinoma after resection. PLoS One 7:e33655.

20. Caruso RA, Bellocco R, Pagano M, Bertoli G, Rigoli L, Inferrera C. 2002. Prognostic value of intratumoral neutrophils in advanced gastric carcinoma in a high-risk area in northern Italy. Mod Pathol 15:831–7.

21. Fu H, Ma Y, Yang M, Zhang C, Huang H, Xia Y, Lu L, Jin W, Cui D. 2016. Persisting and Increasing Neutrophil Infiltration Associates with Gastric Carcinogenesis and E-cadherin Downregulation. Sci Rep 6:29762.

22. Chang HH, Oh PY, Ingber DE, Huang S. 2006. Multistable and multistep dynamics in neutrophil differentiation. BMC Cell Biol 7:11.

23. Su B, Ceponis PJ, Lebel S, Huynh H, Sherman PM. 2003. Helicobacter pylori activates Toll-like receptor 4 expression in gastrointestinal epithelial cells. Infect Immun 71:3496–502.

24. Backert S, Selbach M. 2008. Role of type IV secretion in Helicobacter pylori pathogenesis. Cell Microbiol 10:1573–81.

25. Franchi L, Amer A, Body-Malapel M, Kanneganti TD, Ozoren N, Jagirdar R, Inohara N, Vandenabeele P, Bertin J, Coyle A, Grant EP, Nunez G. 2006. Cytosolic flagellin requires Ipaf for activation of caspase-1 and interleukin 1beta in salmonella-infected macrophages. Nat Immunol 7:576–82.

26. El-Omar EM, Carrington M, Chow WH, McColl KE, Bream JH, Young HA, Herrera J, Lissowska J, Yuan CC, Rothman N, Lanyon G, Martin M, Fraumeni JF, Jr., Rabkin CS. 2001. The role of interleukin-1 polymorphisms in the pathogenesis of gastric cancer. Nature 412:99.

27. Figueiredo C, Machado JC, Pharoah P, Seruca R, Sousa S, Carvalho R, Capelinha AF, Quint W, Caldas C, van Doorn LJ, Carneiro F, Sobrinho-Simoes M. 2002. Helicobacter pylori and interleukin 1 genotyping: an opportunity to identify high-risk individuals for gastric carcinoma. J Natl Cancer Inst 94:1680–7.

28. Peek RM, Jr., Fiske C, Wilson KT. 2010. Role of innate immunity in Helicobacter pylori-induced gastric malignancy. Physiol Rev 90:831–58.

29. Ekman AK, Cardell LO. 2010. The expression and function of Nod-like receptors in neutrophils. Immunology 130:55–63.

30. Jeong YJ, Kang MJ, Lee SJ, Kim CH, Kim JC, Kim TH, Kim DJ, Kim D, Nunez G, Park JH. 2014. Nod2 and Rip2 contribute to innate immune responses in mouse neutrophils. Immunology 143:269–76.

31. Koch M, Mollenkopf HJ, Meyer TF. 2016. Macrophages recognize the Helicobacter pylori type IV secretion system in the absence of toll-like receptor signalling. Cell Microbiol 18:137–47.

32. Miao EA, Andersen-Nissen E, Warren SE, Aderem A. 2007. TLR5 and Ipaf: dual sensors of bacterial flagellin in the innate immune system. Semin Immunopathol 29:275–88.

33. Zhao Y, Shao F. 2015. The NAIP-NLRC4 inflammasome in innate immune detection of bacterial flagellin and type III secretion apparatus. Immunol Rev 265:85–102.

34. Gewirtz AT, Yu Y, Krishna US, Israel DA, Lyons SL, Peek RM, Jr. 2004. Helicobacter pylori flagellin evades toll-like receptor 5-mediated innate immunity. J Infect Dis 189:1914–20.

35. Matusiak M, Van Opdenbosch N, Vande Walle L, Sirard JC, Kanneganti TD, Lamkanfi M. 2015. Flagellin-induced NLRC4 phosphorylation primes the inflammasome for activation by NAIP5. Proc Natl Acad Sci U S A 112:1541–6.

36. Kwok T, Zabler D, Urman S, Rohde M, Hartig R, Wessler S, Misselwitz R, Berger J, Sewald N, Konig W, Backert S. 2007. Helicobacter exploits integrin for type IV secretion and kinase activation. Nature 449:862–6.

37. Forlow SB, Ley K. 2001. Selectin-independent leukocyte rolling and adhesion in mice deficient in E-, P-, and L-selectin and ICAM-1. Am J Physiol Heart Circ Physiol 280:H634–41.

38. Lutz MB, Kukutsch N, Ogilvie AL, Rossner S, Koch F, Romani N, Schuler G. 1999. An advanced culture method for generating large quantities of highly pure dendritic cells from mouse bone marrow. J Immunol Methods 223:77–92. http://doi.org

